# Bi-modal Variational Autoencoders for Metabolite Identification Using Tandem Mass Spectrometry

**DOI:** 10.1101/2021.08.03.454944

**Authors:** Svetlana Kutuzova, Christian Igel, Mads Nielsen, Douglas McCloskey

## Abstract

A grand challenge of analytical chemistry is the identification of unknown molecules based on tandem mass spectrometry (MS/MS) spectra. Current metabolite annotation approaches are often manual or partially automated, and commonly rely on a spectral database to search from or train machine learning classifiers on. Unfortunately, spectral databases are often instrument specific and incomplete due to the limited availability of compound standards or a molecular database, which limits the ability of methods utilizing them to predict novel molecule structures. We describe a generative modeling approach that can leverage the vast amount of unpaired and/or unlabeled molecule structures and MS/MS spectra to learn general rules for synthesizing molecule structures and MS/MS spectra. The approach is based on recent work using semi-supervised deep variational autoencoders to learn joint latent representations of multiple and complex modalities. We show that adding molecule structures with no spectra to the training set improves the prediction quality on spectra from a structure disjoint dataset of new molecules, which is not possible using bi-modal supervised approaches. The described methodology provides a demonstration and framework for how recent advances in semi-supervised machine learning can be applied to overcome bottlenecks in missing annotations and noisy data to tackle unaddressed problems in the life sciences where large volumes of data are available.

## Introduction

Compound identification from tandem mass spectrometry (MS/MS) spectra is one of the grand challenges in analytical chemistry. Automated pipelines support high-throughput projects in various applications, such as clinical diagnostics, natural products discovery and environmental studies ^1–6^. However, with the undeniable progress in the untargeted metabolomics allowing to process large amounts of samples with high resolution and sensitivity^7^, a large share of metabolites in the majority of projects remain unidentified, often referred to as metabolomics dark matter ^8^. The share of identified metabolites is commonly estimated to only be around 2% ^8–10^. Our goal is to develop a scalable machine learning based system for predicting compounds from mass spectra and vice versa that is able to exploit both annotated and unannotated spectra and chemical structures during training.

The availability of high quality and well annotated (i.e., including complete metadata such as instrument, collision energy, etc.) mass spectra is critical to improving compound annotation coverage. Dedicated data collection projects, as well as implementation of FAIR data principles ^11^ in metabolomics, lead to higher increments in the amounts of accessible data. The number of annotated tandem mass spectra in open source libraries (GNPS ^12^, MassBank ^13^, HMDB ^14^, MassBank of North America (MoNA)^1^) is growing every year as efforts of the collective community have generated millions of publicly available spectra. The larger the data volumes are, the more emphasis there is on computational tools capable of handling big data challenges. Current computational approaches to small molecule identification from mass spectra can be generally categorized as the following: 1) spectrum-to-molecule where molecule structure is predicted given a spectrum, or 2) combinatorial in-silico spectra prediction where a spectral database is populated with predicted in-silico spectra derived from molecule structures and then searched for the most likely candidate.

Most of the spectrum-to-molecule methods rely on molecule structural fingerprints ^15–17^, where a multilabel classification problem is solved for predicting if a pre-defined substructure is present, followed by a molecular database search for the most similar fingerprint ^18–20^. Some methods also rely on fragmentation trees ^21^ and spectral trees ^22^ for explicit structural interpretation of spectral data ^18,19,23,24^. The fragmentation tree objects are commonly used for defining kernel functions for kernel-based machine learning methods, e.g. support vector machines ^25^.

Combinatorial in-silico spectra prediction approaches such as CFM-ID ^9^, MetFrag ^26^, FiD ^27^ iterate through possible chemical bonds to identify which cleavage events have more probability to result in the observed set of spectral peaks. Other methods are rule-based and rely on pre-defined physical laws, e.g., ^28^, Mass Frontier™ (ThermoFisher, CA; HighChem, Bratislava, Slovakia).

The choice of a numerical representation of a molecule is a key component of method design. While structural fingerprints have an advantage of being human interpretable, the spaces of most common distance metrics between the fingerprints tend to be ill suited for visualisation and exploration of chemical space ^29^. Samanta et al.^30^ describe numerical representations constructed by deep learning that can overcome these limitations.

Deep neural networks have recently been used to convert from structural fingerprints to spectrum ^31^ and from a spectrum to structural fingerprints ^20^. While demonstrating solid predictive power, the methods still rely on structural fingerprints for the molecule numerical representation, only allowing the deep networks to learn from the predefined molecule substructures. Huber et al.^32^ suggests an adaption of the word2vec algorithm ^33^, which was developed for natural language processing, for mass spectra representation, allowing for more efficient search in the spectral space with no use of information on the molecules structures. To fully utilise the advantages of deep learning approach, we aimed to combine the efficient numerical representations of both spectra and molecules.

Autoencoders ^34^ are a class of generative algorithms for unsupervised machine learning, where a high dimensional input is transformed into a vector of smaller dimension using deep neural networks as an encoder and a decoder. Variational autoencoders (VAEs) ^35,36^ enforce a Gaussian-like distribution of latent space vectors, allowing for easy sampling of realistic looking reconstructions ^37–39^. Multi-modal autoencoders ^40–44^ combine several encoded inputs (modalities) into a coherent joint latent space. Several of them, like SVAE (that was developed for the task of compound identification from mass spectra, thus called SpectraVAE)^44^, allow for missing modalities during training.

We have developed a deep generative approach for furthering efforts towards enabling accurate and robust metabolite identification from MS/MS spectra. The model was based on recent work using semi-supervised deep VAEs to learn joint latent representations of multiple and complex modalities. Conceptually, the model design was based on three main principles: 1) scalability to enable the processing of a growing volume of open access MS/MS spectra; 2) utilisation of both annotated and unannotated spectra and chemical structures; and 3) bidirectional predictions of both spectra to structures and structures to spectra. This functionality was implemented with a multi-modal variational autoencoder model where the modalities included MS/MS spectra and molecular structures.

In this study, we define a paired sample as an MS/MS spectrum with an annotated molecule structure; An unpaired sample was considered an MS/MS spectrum without an annotated molecule structure or a molecule structure without an MS/MS spectrum. The SVAE model allowed for constructing a joint latent representation of MS/MS spectra and molecules structures using paired and unpaired samples. We hypothesized that the millions of unpaired samples available in MS/MS spectra and molecular structures in publicly available databases and datasets could be leveraged to improve the quality of the joint latent space, thus improving the ability of the model to reconstruct MS/MS spectrum and molecules structures even with missing input data (i.e., either the MS/MS spectrum or the molecule structure).

To the best of our knowledge, the presented model is the first to apply semi-supervised generative learning to the compound identification problem, as well as the first algorithm to bidirectionally make predictions of molecule structures given the spectra and of spectra given the molecule structures. We compared SVAE to another bi-modal VAE-based model, proving SVAE’s better predictive power. We studied the effect of adding unpaired molecules to the training set and demonstrated the superiority of the semi-supervised approach. Different numerical representations of molecules were analysed, one-hot encoded SMILES strings and structural fingerprints. The structural fingerprints models were benchmarked against other methods on Critical Assessment of Small Molecule Identification (CASMI2017) ^45^ dataset.

## Methods

### Multi-modal variational autoencoders

A variational autoencoder (VAE) ^35^ is an unsupervised machine learning algorithm, where a latent representation *z* of an observed variable *x* is generated using the model *p*(*x,z*) = *p*(*z*)*p*(*x|z*). The intractable posterior *q*(*z|x*) and the conditional distribution *p*(*x|z*) are approximated by neural networks using the loss function:

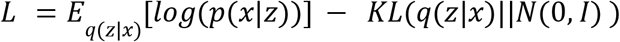

where *KL* is Kullback-Leibler divergence, a measure of how similar the distributions are, and *N*(0,*I*) is a standard normal distribution.

For the purpose of multi-modal learning, the variational autoencoder framework is extended to have an encoder and decoder network for each modality. The main problem is how to enforce the latent representations derived for each modality to be coherent, so one modality can be used to reconstruct another. In ^44^ we derived an SVAE model, which takes the form:

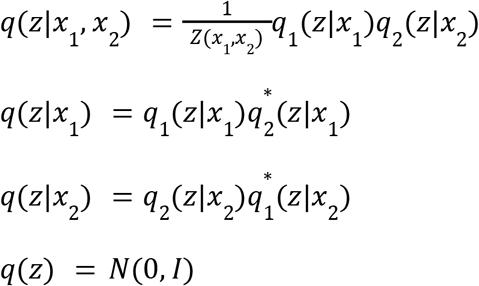

The distributions *q*_1_(*z|x*_1_), *q*_2_(*z|x*_2_), 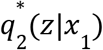 and 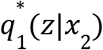 are approximated by neural networks. See Supplementaries A for the full definition of ELBO-type loss for this model.

In the current study we compared SVAE to JMVAE ^40^, the approach that scored among the top for fully supervised (all the samples are paired) bi-modal computer vision datasets. The JMVAE model approximates *q*(*z|x*_1_, *x*_2_), *q*_1_(*z|x*_1_) and *q*_2_(*z|x*_2_) with three neural networks and optimizes an ELBO-type loss of the form

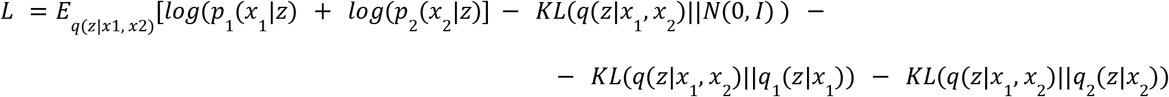

A graphical explanation for SVAE and JMVAE models is provided in Figure 1.

**Figure 1.**
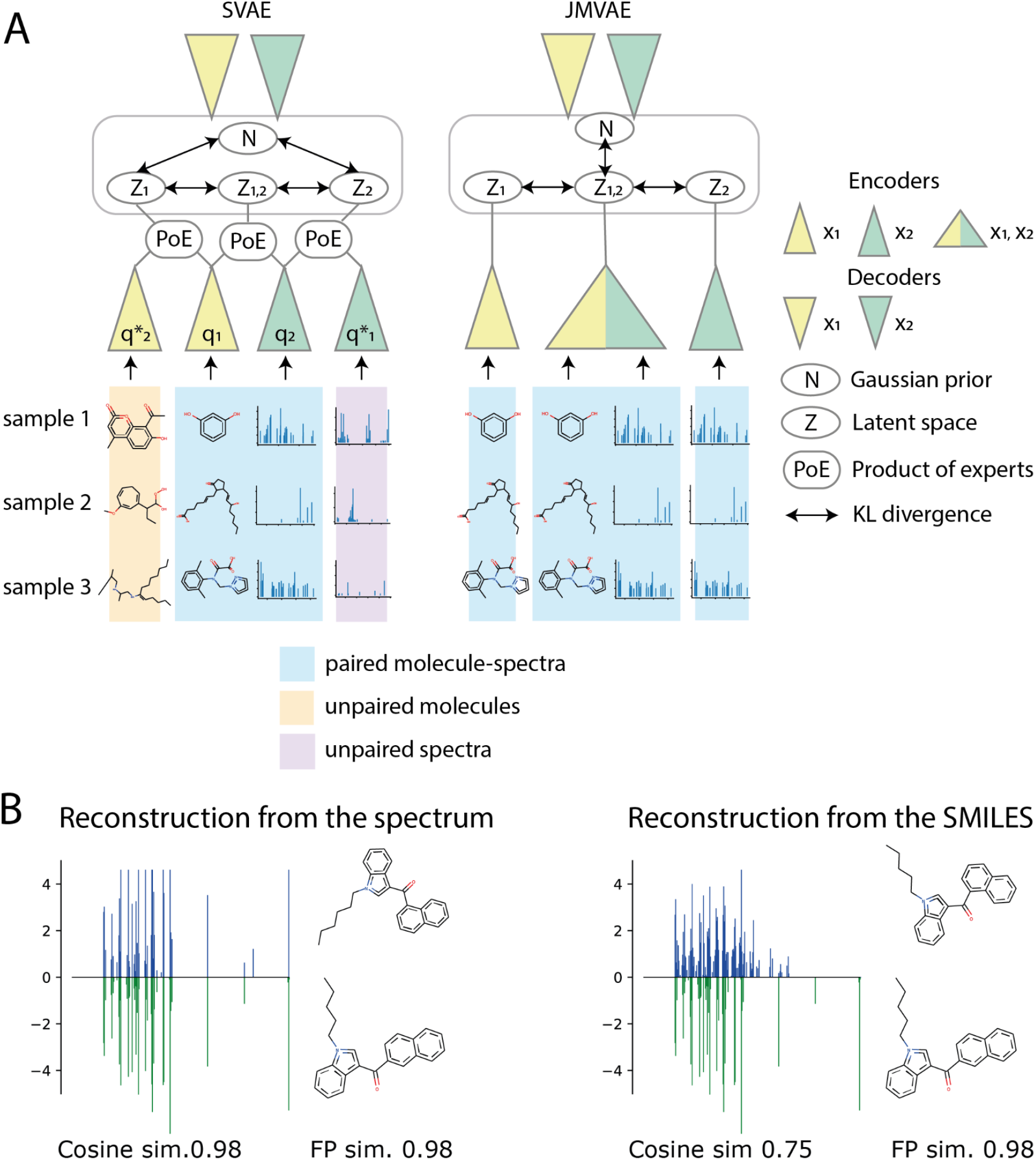
(A) Schematic architecture overview of compared bi-modal VAEs. SVAE ^44^ can process both paired and unpaired samples, allowing for semi-supervised learning. JMVAE ^40^, can only process paired samples. Each triangle stands for an individual neural network, the network colors indicate the different modalities. (B) An example of input and output data during testing. The bottom part of reconstructions shows input data, a spectrum (in green) and a molecule. The upper part shows the reconstructions for a spectrum (in blue) and a molecule. (left) Example reconstructions where only the spectrum was provided as an input; (right) Example reconstructions where only the molecule was provided as an input. Cosine similarity was calculated between input and output spectra, and fingerprint similarity was calculated between input and output molecules.

In ^44^ we also benchmarked VAEVAE against the SVAE model on the metabolite identification task with the supervised dataset. SVAE performed better, so we did not include VAEVAE in an evaluation used in this paper.

### Dataset

Following Fan et el.^20^ we used the same homogeneous tandem mass spectra dataset. The spectra were acquired from Mass Bank of North America (MoNA)^2^ and NIST^3^ libraries. The selected spectra are recorded in positive ion mode with collision cell or ion trap instruments with a mass range from 100 to 1100 Da. The full list of spectra IDs can be found at https://github.com/sgalkina/svae_spectra. Spectra were binned with the bin size 0.1Da, intensities of the peaks that ended up in the same bin were summed. The resulting vector *v* of length 11000 was further processed using the function *v_l_* = *log*(1 + *v*). *v_l_* was an input to the spectral modality encoder.

A common strategy for building datasets for molecular structure predictions is to sum all the spectra collected for the same structure (InChiKey) to one. Our dataset was assembled from both summed and original spectra.

The unpaired molecule structure dataset consisted of around 2 million biomolecules, a subset of which was used as one of the built-in databases by SIRIUS software ^18^. The biomolecules were from similar compound classes as the spectral dataset. The fingerprints for the spectra-fingerprints experiment (the section ”Spectra and structural fingerprints”) were calculated for the same dataset. The full list of SMILES can be found at https://github.com/sgalkina/svae_spectra. Unpaired spectra were not used in this study, see the explanation in Supplementaries C.

### Encoders and decoders architectures

We showed two approaches for metabolite identification: reconstructing a SMILES string (Section ”Spectra and SMILES”) and reconstructing a molecule structural fingerprint (Section ”Spectra and structural fingerprints”). The detailed description of all the network architectures is in Supplementaries E.

Both approaches used the same encoder and decoder networks for the spectrum modality. The spectrum encoder consisted of several fully connected layers with batch normalisation. The decoder was symmetrical to the encoder. See also Supplementaries C for an alternative architecture that used TreeLSTM ^46^ to process fragmentation trees ^21^.

Constructing meaningful latent representations of a molecule structure using the VAE framework is ongoing research ^47–51^. In this study, we used a string-based model ^52^ that works directly with SMILES strings. The latent space constructed by this model was also tested as a molecule similarity metric ^30^.

For the structural fingerprints encoder and decoder we used symmetrical networks consisting of 4 fully connected layers.

## Results

To demonstrate the effect of training a bi-modal VAE on unpaired molecules we compared SVAE trained on paired and unpaired data to SVAE trained only on paired data. As a baseline, we considered the JMVAE ^40^ model which can only be trained on paired data and showed good results in this setting when compared to other bi-modal VAEs in label prediction tasks ^42^.

To benchmark bi-modal VAEs against the current metabolite identification methods and against a supervised multilabel classification model, we used structural fingerprints as a second modality and evaluated the results on the CASMI challenge.

To evaluate the quality of reconstructed spectra, we used the SIRIUS algorithm as an independent oracle (see Section “Predicting spectra”).

### Spectra and SMILES

The bi-modal variational autoencoder model design allowed for missing inputs at test time. We tested the quality of four reconstructions: SMILES to SMILES, spectrum to spectrum, SMILES to spectrum, and spectrum to SMILES. A SMILES to SMILES and a spectrum to spectrum reconstruction tested the model capacity for building latent representations for an individual modality. A spectrum to SMILES and a SMILES to spectrum tested the predictive power across the two modalities.

Here, semi-supervised SVAE represented the SVAE model trained on both 36 thousand paired samples and 2 million unpaired molecules. Supervised SVAE represented the SVAE model only trained on the paired samples. The JMVAE training set was the same as for supervised SVAE.

The evaluations were performed using a structure-disjoint test set of 1000 spectra for 950 unique molecules (772 unique InChiKeys). The molecules were not a part of the unpaired molecule training set.

The aggregation of quantitative results is presented in the Tables 1–4 and the Figure 3. The source code can be found at https://github.com/sgalkina/svae_spectra.

**Table 1.** Quantitative evaluation of molecule reconstructions given a spectrum and a SMILES string on a test set of 1000 molecules. Percentage of valid SMILES is calculated for 1000 attempts to reconstruct a molecule. Fingerprint similarity is Tanimoto distance between the Daylight fingerprints calculated with RDKit, the metrics is from 0 to 1 with 1 showing the most similarity. Other metrics are calculated as Maximum Mean Discrepancy (MMD) between input and output molecule graphs distributions: graph degree (Degree), clustering coefficient (Cluster) and an average orbit counts statistics, the number of occurrences of all orbits with k nodes (Orbit). The lower the distance is, the more similar the distributions are.

From the spectrum
|  | % of valid<br>SMILES | Fingerprint<br>(mean / std) | similarity | Degree | Cluster | Orbit |
| --- | --- | --- | --- | --- | --- | --- |
| JMVAE | 99.6 | 0.096/0.095 |  | 0.140 | 0.0003 | 0.011 |
| SVAE, supervised | 97.5 | 0.131/0.116 |  | 0.125 | 0.0003 | 0.009 |
| SVAE, semi-supervised | 95.2 | 0.185/0.160 |  | 0.059 | 2.716-05 | 0.005 |
| Baseline, trivial molecules | 100 | 0.044/0.043 |  | 0.235 | 0.0005 | 0.013 |

**Table 2.**
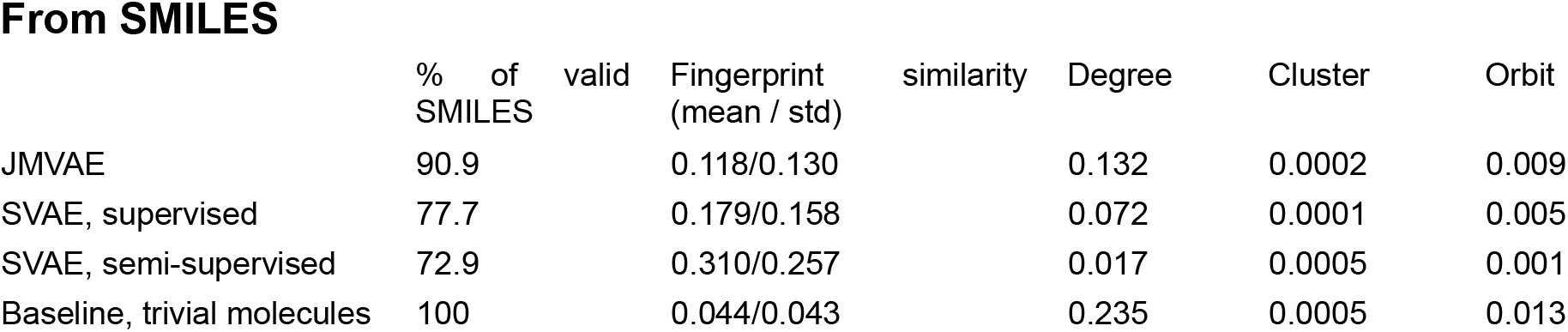
Quantitative evaluation of molecule reconstructions given a spectrum and a SMILES string on a test set of 1000 molecules. Percentage of valid SMILES is calculated for 1000 attempts to reconstruct a molecule. Fingerprint similarity is Tanimoto distance between the Daylight fingerprints calculated with RDKit, the metrics is from 0 to 1 with 1 showing the most similarity. Other metrics are calculated as Maximum Mean Discrepancy (MMD) between input and output molecule graphs distributions: graph degree (Degree), clustering coefficient (Cluster) and an average orbit counts statistics, the number of occurrences of all orbits with k nodes (Orbit). The lower the distance is, the more similar the distributions are.

**Table 3.**
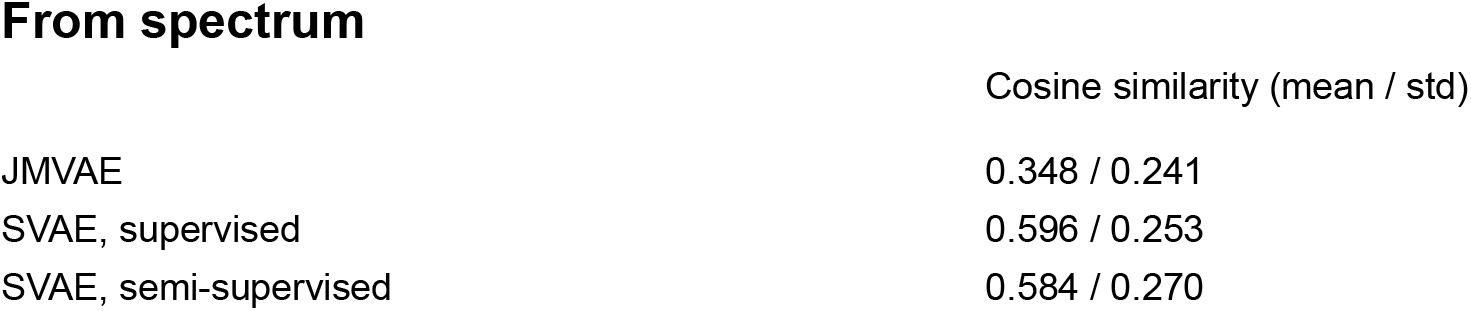
Quantitative evaluation of spectra reconstructions given a spectrum and a SMILES string on a test set of 1000 molecules. Because all the spectra vectors are positive, cosine similarity here is a metrics from 0 to 1 with 1 showing the most similar spectra.

**Table 4.**
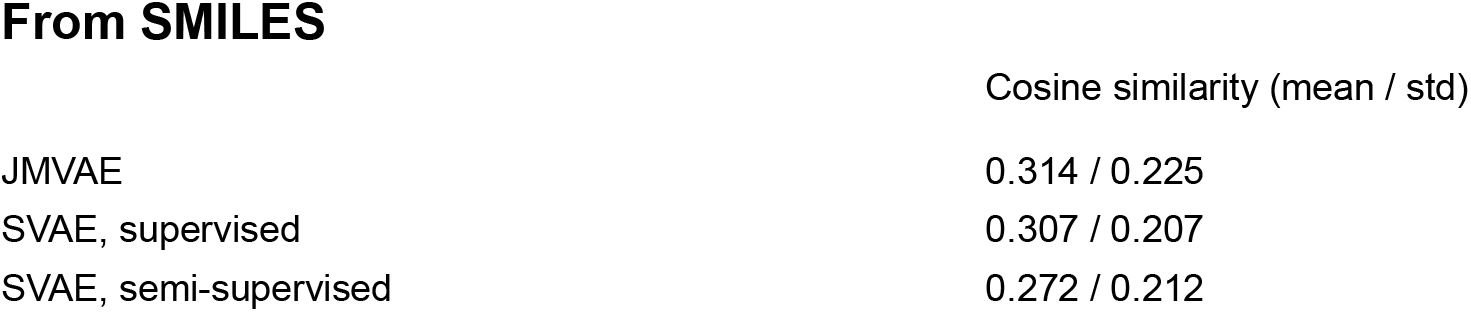
Quantitative evaluation of spectra reconstructions given a spectrum and a SMILES string on a test set of 1000 molecules. Because all the spectra vectors are positive, cosine similarity here is a metrics from 0 to 1 with 1 showing the most similar spectra.

From SMILES
|  | Cosine similarity (mean / std) |
| --- | --- |
| JMVAE | 0.314 / 0.225 |
| SVAE, supervised | 0.307 / 0.207 |
| SVAE, semi-supervised | 0.272 / 0.212 |

#### Predicting molecules

Following Gomez-Bombarelli et al.^52^ we used 1000 attempts to generate a valid molecule from a test set sample. The percentage of the resulting valid molecules was one of the evaluation metrics. Each reconstructed molecule was compared to the corresponding original molecule by RDKit topological fingerprint similarity. Additionally, the set of reconstructed molecules was compared with the set of original molecules by its degree, cluster and orbit similarity metrics ^48^ for the reconstruction of a wide range of graphs.

It is important to notice that during the early epochs of training the output of the network molecule decoder is a string of C letters of the length similar to the length of the original SMILES. The same kind of output was observed in cases when a trained network failed to reconstruct a molecule (see the reconstructions examples in the Supplementaries). These reconstructions, however non-informative, are valid SMILES strings and therefore contribute to the overall percentage of valid reconstructions. It explains why the reconstruction from the spectrum results in a higher share of valid molecules for all the models than a less complicated task of reconstructing a SMILES string given a SMILES string. We called the strings of C letters of the same length as the original SMILES a “trivial molecule” and include the evaluation of the trivial molecules set as a baseline for all the metrics in the Tables 1–2.

The evaluation results for the basic variational autoencoder for SMILES string with the same SMILES encoder and decoder architecture as the bi-modal network is shown in Figure 3. The quality of the individual modality reconstruction became worse than from a one-modality VAE, since performing cross-modality learning and bi-directional reconstructions may interfere with more specialised one modality reconstruction learning.

#### Predicting spectra

Cosine similarity is a common metric for quantitative evaluation of spectra similarity. While it is a meaningful metric for the spectra-to-spectra reconstruction, it does not capture the quality of SMILES-to-spectra reconstruction for our models. The models were trained on the tandem mass spectra of the same ionisation mode, but different collision energies, including the spectra for which all the available collision energies were merged into one spectrum. Besides the fact that a probabilistic model will have variance in the output, the current model is not conditioned on a specific collision energy, which means that the spectra reconstructed from SMILES strings form a distribution over all the collision energies the model is trained on.

To overcome the limitations of cosine similarity metrics for SMILES-to-spectra reconstruction, we evaluated how well the reconstructed spectra can predict molecular structures using SIRIUS software as an independent oracle. 100 spectra were independently reconstructed for each of the random 100 SMILES strings from the structure disjoint test set. Each of these spectra was given to both the same SVAE model and the SIRIUS software as an input. SIRIUS also requires a precursor m/z value as an input, so the corresponding value associated with the original data entry was added. One candidate molecule was predicted by SIRIUS and SVAE per each spectrum, resulting in 100 candidate molecules per one input molecule. For SIRIUS, the molecule is the top candidate, and for SVAE it is the first valid reconstruction from 1000 attempts. Mean fingerprint similarity between all the candidates for each input molecule is reported in Figure 2D-1, and the distribution of corresponding variances for the SIRIUS algorithm is in Figure 2D-2.

**Figure 2.**
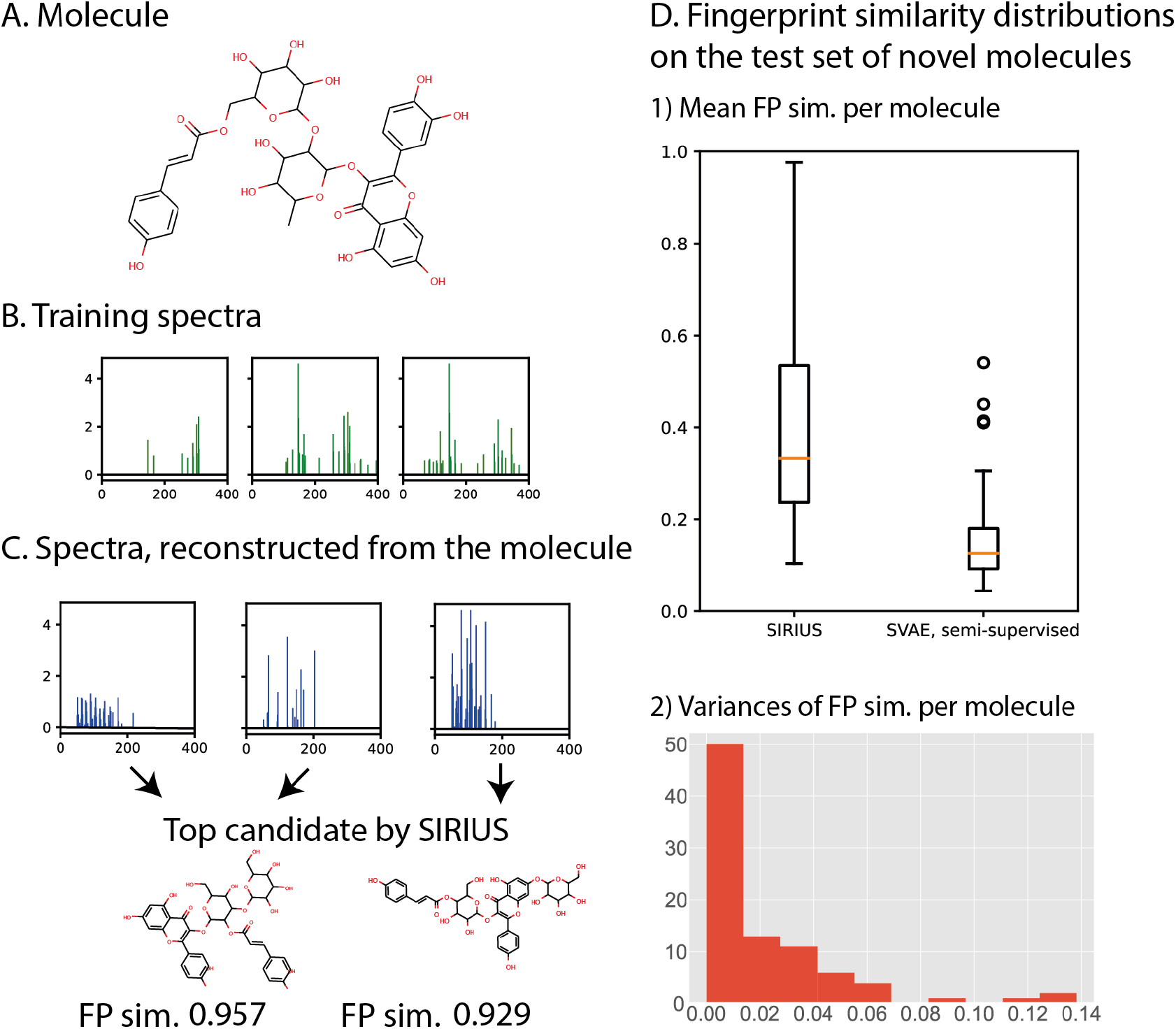
A. A molecule from the training set. B. 3 examples of training spectra for this molecule. The training spectra for this molecule have different collision energies ranging from 10 to 50eV. C. 3 independent reconstructions of the spectrum given the same unpaired molecule as input. Because the training set spectra were different, the generated reconstructed spectra may differ as well resulting in low values for the similarity metrics between one original spectrum and the reconstruction from the corresponding SMILES. To evaluate the spectra with an independent oracle method, we predicted molecule structures for the generated spectra using SIRIUS. The top candidates do not always match the exact molecule structure, but have structural resemblance. The number below the molecules indicate the fingerprint similarity (FP sim.) between an original molecule and the SIRIUS top candidate molecule for the predicted spectrum. D. 1) The distribution of fingerprint similarities of the molecules predicted by the reconstructed spectra for a test set of 100 molecules. The test set consisted of structure-disjoint molecules. Each point on the distribution represents the fingerprint similarity averaged across molecule predictions for 10 spectra. If a spectrum did not have hits by the SIRIUS software, it was not included in the statistics. “SIRIUS top candidate” distribution provides predictions made by the SIRIUS software, and “SVAE semi-supervised” provides predictions of molecules made by the semi-supervised SVAE model from the reconstructed spectra. 2) The distribution of variances for fingerprint similarities of 100 candidates per each input molecule. The variance histogram and the selected number of candidates suggest a reliable estimate of a mean fingerprint similarly per input molecule.

For 95% of input molecules the standard deviation of a fingerprint similarity estimate does not exceed 0.25 (Figure 2D-2). With the sample size of 100 for reconstructed spectra the standard deviation for mean fingerprint similarity does not exceed 0.025, making the distribution of sample means (Figure 2D-1) a reliable estimate.

#### CASMI2017

We evaluated the models on the CASMI2017 challenge. Since all the training spectra were collected in positive ion mode, we only considered the positive spectra for the evaluation. All the predicted molecule structure were ranked by the fingerprint similarity with the original molecule, 10 predicted molecule structures with the highest fingerprint similarity are shown on Figure 4A. The predicted molecules with lower scores than top 10 resemble “trivial” molecules more than original ones.

**Figure 3.**
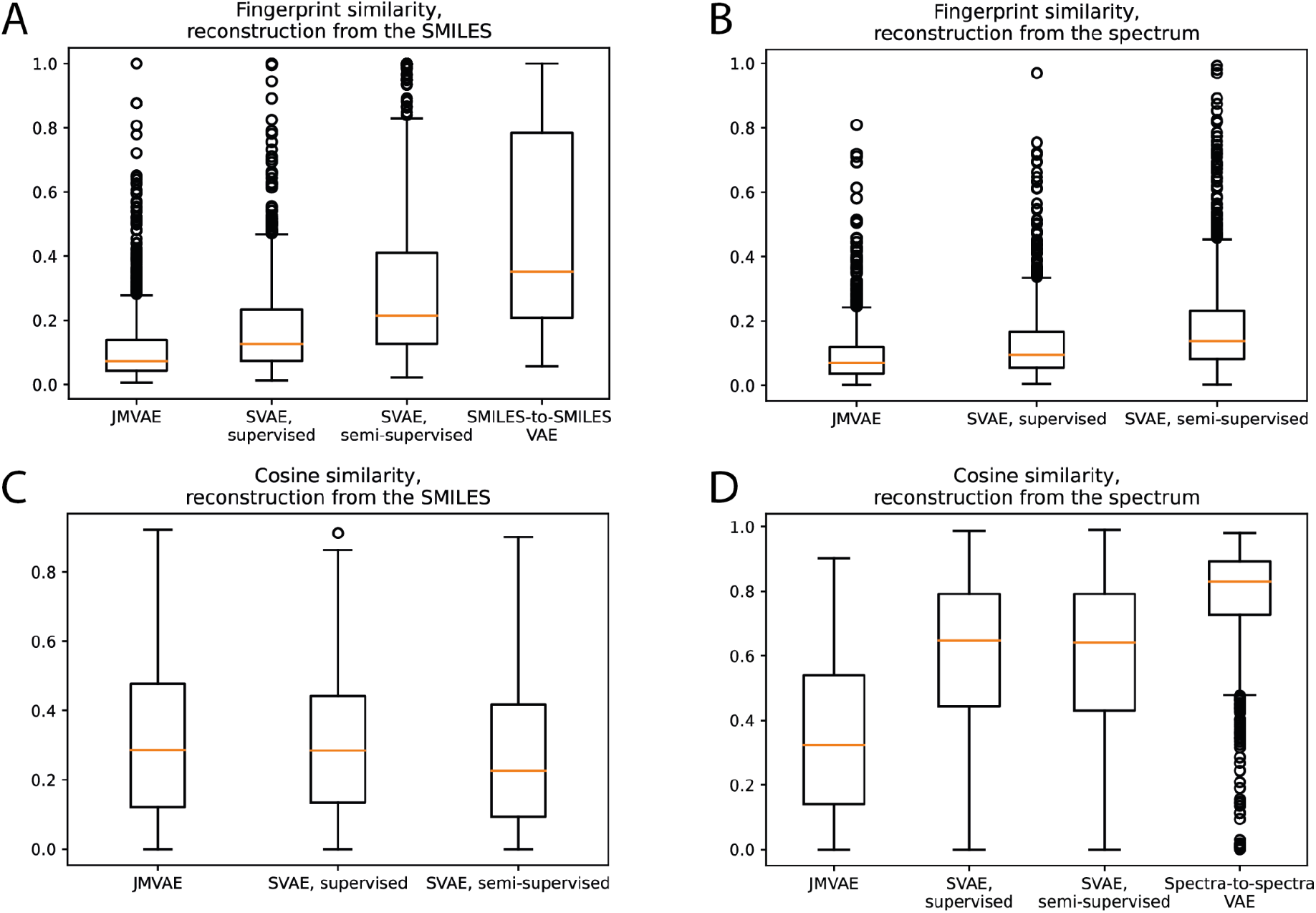
Distributions of similarity metrics for each possible reconstruction. A. Distributions of fingerprint similarities between an original molecule and a reconstruction from the SMILES string. Semi-supervised SVAE shows the strongest performance. Basic SMILES-to-SMILES variational autoencoder trained on 2mil biomolecules (same as semi-supervised SVAE, but no spectral modality) results are presented for the reference and show that bi-direction learning decreases the quality of one modality reconstruction. B. Distributions of fingerprints similarities between an original molecule and a reconstruction from the spectrum. The performance ranking between the models was the same as for the SMILES-to-SMILES reconstruction. C. Distributions of cosine similarities between an original spectrum and a reconstruction from the SMILES. This metric was not the most optimal for the current composition of the training set as discussed in the Section “Predicting spectra”. D. Distributions of cosine similarities between an original spectrum and a reconstruction from the spectrum. Since both supervised and semi-supervised SVAE were trained on the same spectral dataset, both SVAE models show similar results, outperforming the JMVAE model.

**Figure 4.**
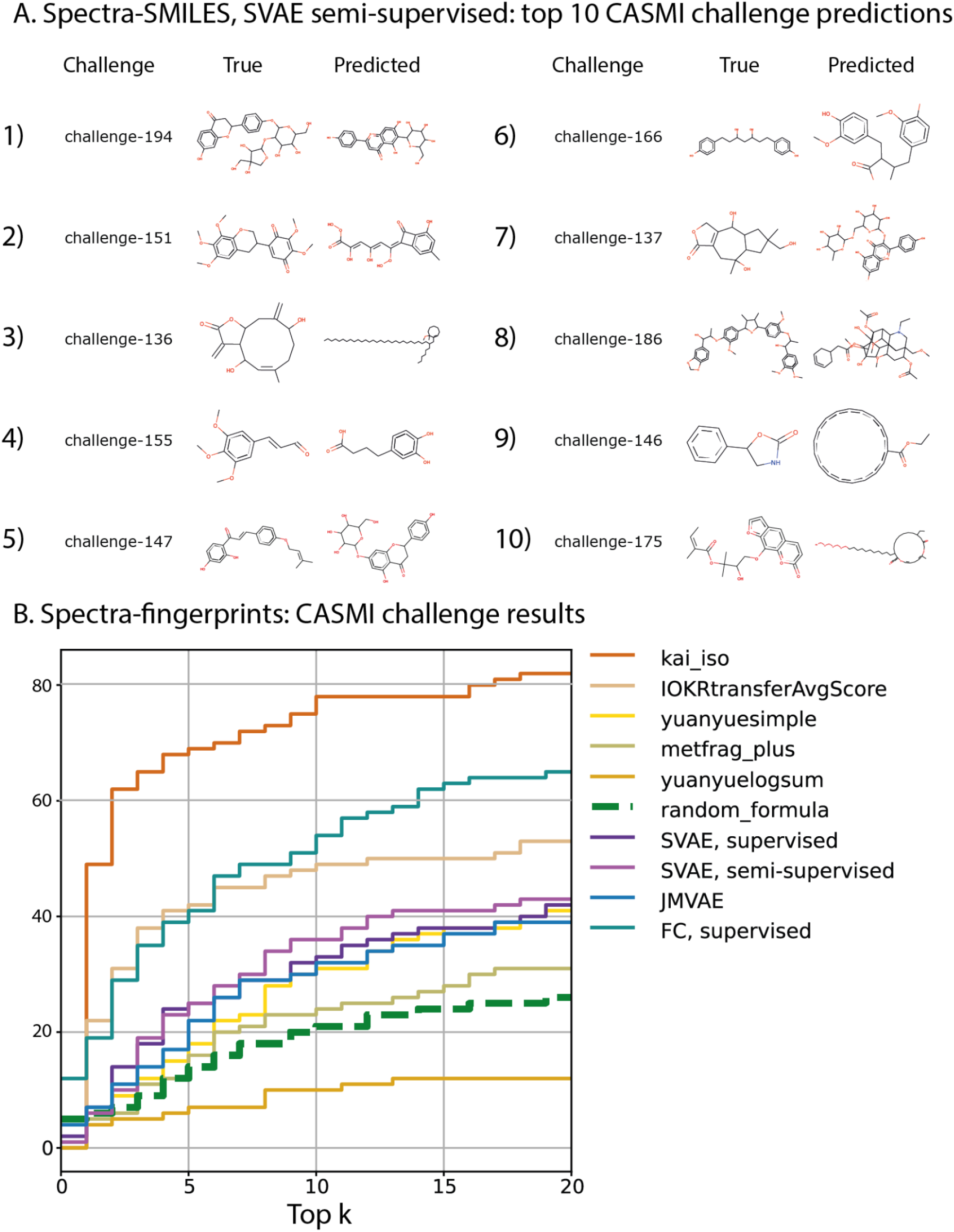
A. CASMI2017, positive ion mode challenges, top 10 molecule reconstructions from the spectrum sorted by the fingerprint similarity between the true molecule and the molecule, reconstructed from the spectrum. The molecules with the reconstruction quality below the top 10 resemble trivial molecules and do not have meaningful structural information. B. CASMI2017 leaderboard with fingerprints. The x-axis represents the number of top candidates, the y-axis represents how many of 112 positive ion mode spectra the correct molecule was among top *k* candidates. kai_iso, IOKRtransferAvgScore, yuanyuesimple, metfrag_plus and yuanyuelogsum were the competing algorithms. FC, supervised was a deep learning network that predicts a structural fingerprint from a spectrum. SVAE, semi-supervised, SVAE, supervised and JMVAE were the bi-modal VAEs.

**Figure 5.**
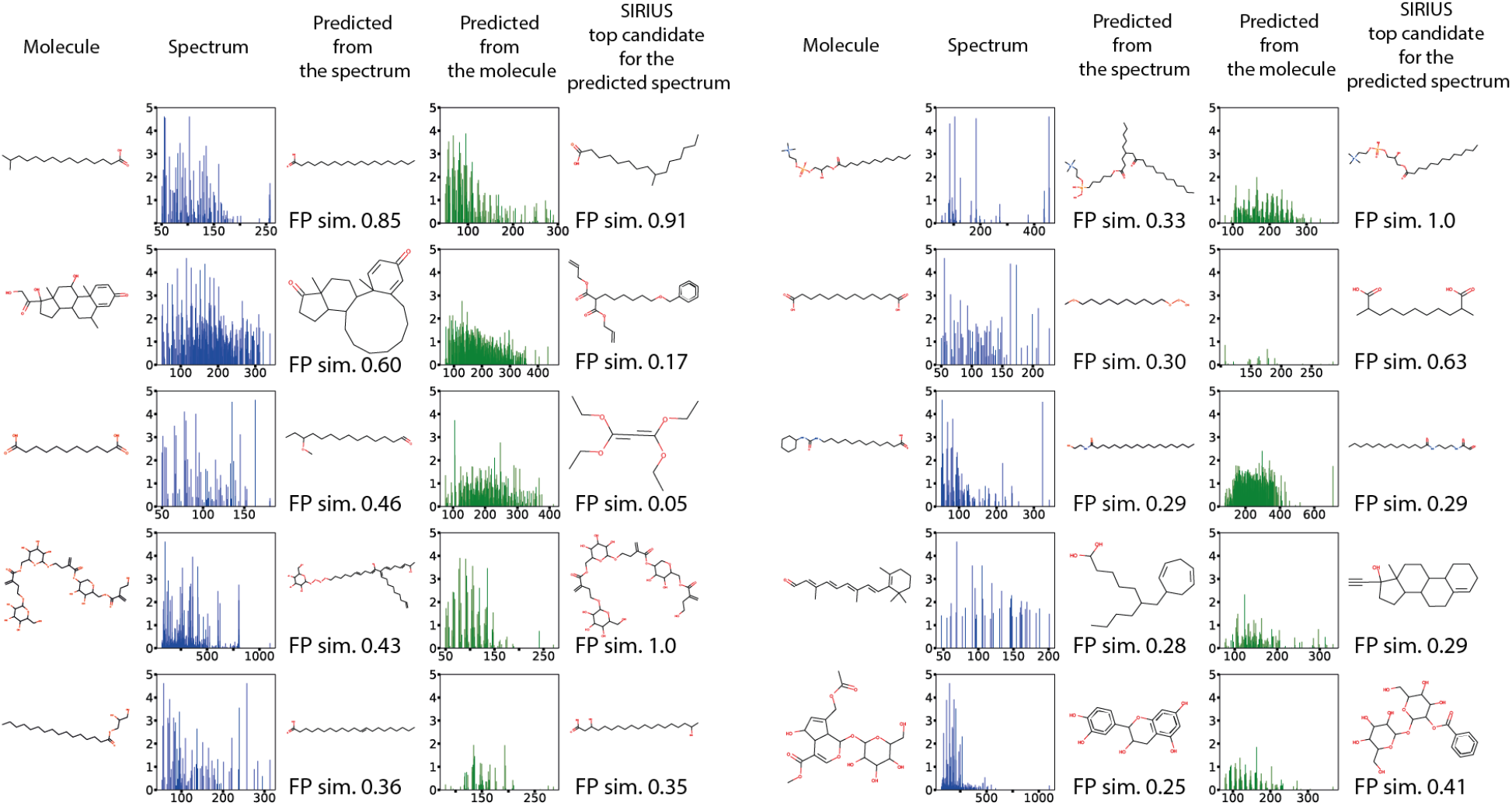
Examples of lipid molecules and spectra reconstructions from semi-supervised SVAE trained with no lipid-like spectra.

The setup of the CASMI2017 challenge was that a method must provide a scored list of candidate molecules, with a higher non-negative score indicating the better candidate. Our method was not able to provide the scoring for the given list of candidates because our method did not involve a molecule database search. To benchmark bi-modal variational autoencoders against other methods we evaluated it on a different representation of a molecule, a structural fingerprint, see the Section ”Spectra and structural fingerprints”.

### Spectra and structural fingerprints

A one-hot encoded SMILES string is not the only way to numerically represent a molecule structure. A different numerical representation requires different encoder and decoder architectures, which can be easily integrated to the bi-modal VAE network. One of the most common ways to numerically represent molecules is via structural fingerprints, which are a bit string where each bit stands for the presence of a predefined molecule substructure ^15,16^. A combination of several fingerprints ^53^ has proven to be an efficient instrument for the molecule identification task.

Often, the molecule identification workflow ^18,20^ includes steps for predicting structural fingerprints and searching for the best fingerprint match in the structure database. The quality of the molecule structure prediction thus depended on the size of the candidates molecular database. For example, the latest published evaluation of SIRIUS algorithm showed that when searching in the full PubChem database the correct identification rate for a test set was 40.4%, but after narrowing the search to only the biomolecule set of 0.5 million molecules, the rate was 74.0% ^18^.

In order to evaluate the quality of fingerprint predictions, we chose the CASMI challenge category with the candidate structures provided. Here, a method returned a sorted list of candidates. The scoring was performed by the cross entropy loss between the predicted fingerprint and the candidate fingerprint. The rank of the correct molecule was its place in the sorted list. We showed for how many of the 112 positive mode spectra the correct candidate has the rank of *k*, for 0 ≤ *k* ≤ 20in Figure 4B. The top performing algorithm is shown for each participating team.

Duhrkop et al.^*18*^ demonstrated that the chemical formula can be predicted from a tandem mass spectrum with 93.8% accuracy. Following Fan et al.^20^, we assumed the molecule formula was known for each spectrum. This allowed us to filter out candidates with the incorrect molecule formula. The remaining candidates were ranked by log loss between their fingerprint and the predicted one. As a baseline method, we picked a random candidate for each spectrum among the candidates with the same molecule formula (shown in Figure 4B as *random_formula*).

We also tested a simpler feed forward deep learning network (*FC, supervised* on Figure 4B). The architecture conceptually resembles Fan et al.^20^, though it was not exactly the same in terms of features, the number and size of hidden layers and which structural fingerprint was predicted (see Supplementaries E for the details). *FC, supervised* had the 2nd best result. The bi-modal VAEs performed worse, returning the results comparable to the 3rd best method. It showed that if a modality is a simple object, like a bit vector in this case, it is more beneficial to train a specialised multilabel classifier instead of a bi-directional VAE: the bi-direction VAE is more useful in applications involving complex and noisy modalities (an observation supported by Wu et al.^41^, where a supervised classifier performed better for predicting the label for a labeled image dataset than a bi-modal VAE with the label as one of the modalities). Semi-supervised SVAE showed better performance compared with the supervised VAE models.

## Discussion

We developed a semi-supervised machine learning approach for the metabolite identification task. In order to avoid using molecular databases to search for the molecule candidates we used a molecule structure generative model. It is integrated with a bi-modal VAE-based model and designed to perform training on both paired and unpaired samples of molecules and MS/MS spectra. The bi-modal setup also allowed for the generation of in-silico mass spectra from molecule structures. To the best of our knowledge, the suggested method was the first to perform bi-directional predictions.

The test set molecules are selected to be structure-disjoint (“novel” for the algorithm). Benchmarking of two different bi-modal VAE architectures (SVAE vs JMVAE) showed that SVAE performs better on a fully paired dataset. SVAE was also trained in a semi-supervised setting, with 2 million molecules from similar compound classes, and performed better than supervised SVAE.

To benchmark against other algorithms, we used a different numerical representation of a molecule, a structure fingerprint. The results for the CASMI challenge data suggest that bi-modal VAEs with the basic architectures perform far from the state-of-the-art (3rd best result), while a more specialised multiclass classification neural network shows the 2nd best result, highlighting the need for future model architecture optimization. The suggested bi-modal framework allows for easy testing of the different encoder architectures.

A major challenge in using publicly available MS/MS spectra databases to annotate unknown MS/MS spectra or infer similar MS/MS spectra is the large heterogeneity of MS/MS spectra derived from the same compound. The heterogeneity in MS/MS spectra is due to the influence of the sample (i.e., sample matrix and solvent), instrumentation factors (e.g., source design, collision cell, detector technology, etc.), and instrument parameters (e.g., carrier gases, collision energy, declustering potential, acquisition rate, etc.) In the work presented, training data was limited to positive ion mode and a single type of instrument, but we did not control for instrument parameters which meant that every compound could have a different set of paired MS/MS spectra. The use of multiple MS/MS spectra acquired using different instrument parameters for each annotated molecule structure improves the diversity of training data, which has been shown to be beneficial to training deep learning models ^54^, but complicates the validation of predicted MS/MS spectra because the generation of MS/MS spectra follows a probability distribution of likely spectra over a range of different instrument parameters. We overcame the challenge of annotating generated MS/MS spectra through the use of an independent Oracle (i.e., Sirius).

Future work will address the ability to explicitly include sample, instrumentation factors, and instrument parameters as input and output. In the case of a current dataset, it would mean including instrument parameters (e.g., collision energy) as an additional input to stylize the spectra accordingly. With the current SVAE architecture, instrument parameters as input can be a third modality. SVAE successfully scales to three modalities ^44^. Another approach would be to disentangle the factors that influence the generation of the MS/MS spectra in the latent space directly. Previous works ^35,55,56^ describe various VAE approaches to condition and disentangle VAE latent representations into independent discrete and continuous factors. Integrating disentangled representations into a multi-modal semi-supervised VAE would allow to generate customised in-silico spectra and will likely lead to better hidden representation and better predictive performance.

Performance of the presented model depends on many factors including the encoder and decoder architectures, loss functions, and input and output representations. In this work, we sought to demonstrate the feasibility and utility of multi-modal and semi-supervised learning leveraging unannotated MS/MS spectra and molecular structures to improve compound identification accuracy from MS/MS experiments. Future work will seek to optimize and tune the model itself. Current deep learning methods that predict information from the mass spectra use binned spectra and process them with multilayer perceptrons ^20,31^, word2vec algorithm inspired by natural language processing ^32^ or a transformer architecture ^57^. A binned spectrum has an obvious drawback of reducing resolution of the original spectrum, losing sensitivity provided by the latest generation of mass spectrometers. An architecture that is able to utilise the exact m/z values and a varied input size, f.e. a recurrent network or another NLP-inspired algorithm, can be easily integrated to the presented bi-modal network. Another possible optimization is integrating fragmentation or spectral trees to the spectra encoder architecture. See Supplementaries C describing a possible use of TreeLSTM ^58^ networks for adding fragmentation trees.

The same applies to the molecule structure encoding task. Numerically encoding a molecule structure is a subject of ongoing research. The relevant methods can be roughly divided into string based ^52,59^ and graph based ^47,48,60–63^. State-of-the-art methods generate a very high share of valid molecules or are designed to generate only valid molecules (e.g. JTNN). Unlike the SMILES string based method, graph based algorithms can parse and process the underlying semantic structure of a molecule graph object. Graph based methods demonstrated an ability to predict molecule properties from the latent space, so integrating such a method to the bi-modal framework will be beneficial.

## Conclusion

We proposed a bi-modal variational autoencoder model capable of learning from data with missing annotations. The developed method was the first demonstration of semi-supervised learning applied towards metabolite identification from MS/MS spectra. The developed model was also the first bi-directional prediction tool to predict molecule structures from MS/MS spectra and MS/MS spectra from molecular structures. We demonstrated the predictive power of the algorithm with multiple experiments that included well established benchmark datasets. The resulting numerical representation of molecules and spectra as encoded in the latent space can also be used in different applications, e.g. molecule property predictions. Future work would also include optimizing encoder and decoder architectures and learning disentangled latent representations of the spectra and molecule objects for better tuning of the in-silico spectra generation process. The developed model provides a general framework for tackling life science problems where large volumes of unannotated, complex, and noisy data are available for training.

## Author Contributions

S.K. and D.M. designed the experiments, S.K., C.I and M.N designed the methods. S.K. wrote the code and performed the experiments. S.K. and D.M. wrote the manuscript, all the authors reviewed the manuscript.

## Footnotes

1 https://mona.fiehnlab.ucdavis.edu/

2 https://mona.fiehnlab.ucdavis.edu/

3 https://chemdata.nist.gov/

