## Supplementaries for "Bi-modal Variational Autoencoders for Metabolite Identification Using Tandem Mass Spectrometry"

### A ELBO-type loss for SVAE

$$
\begin{aligned}
&= E_{p_{\text{paired}}(x_1,x_2)}\left[E_{q(z|x_1,x_2)}[\log p_1(x_1|z) + \log p_2(x_2|z)]\right] \\
&\quad - D_{\text{KL}}(q(z|x_1,x_2) \parallel p(z|x_1)) - D_{\text{KL}}(q(z|x_1,x_2) \parallel p(z|x_2)) \\
&\quad + E_{p_{\text{paired}}(x_1)}\left[E_{q(z|x_1)}[\log p_1(x_1|z)] - D_{\text{KL}}(q(z|x_1) \parallel p(z))\right] \\
&\quad + E_{p_{\text{paired}}(x_2)}\left[E_{q(z|x_2)}[\log p_2(x_2|z)] - D_{\text{KL}}(q(z|x_2) \parallel p(z))\right] \qquad (1)\\
\mathcal{L}_1 &= E_{p_{\text{unpaired}}(x_1)}\left[E_{q(z|x_1)}[\log p_1(x_1|z)] - D_{\text{KL}}(q(z|x_1) \parallel p(z))\right] \qquad (2)\\
\mathcal{L}_2 &= E_{p_{\text{unpaired}}(x_2)}\left[E_{q(z|x_2)}[\log p_2(x_2|z)] - D_{\text{KL}}(q(z|x_2) \parallel p(z))\right] \qquad (3)\\
&= + \mathcal{L}_1 + \mathcal{L}_2 \qquad (4)
\end{aligned}
$$

Here $p_{\text{paired}}$ and $p_{\text{unpaired}}$ denote the distributions of the paired and unpaired training data, respectively. See (Kutuzova et al., 2021) for the full model derivation.

### B Novel molecule class

We evaluated the models ability to generalise to predict spectra and structures for a molecule class that did not have representative spectra in the training set. We selected two molecules classes identified with the ClassyFire tool (Feunang et al. 2016), “Lipids and lipid-like molecules” and “Benzenoids”. For each of those classes, we performed an experiment where we removed the corresponding molecules from the paired spectra-molecules training set. We did not change the unpaired molecules training set, which still contained molecules from this class. We trained semi-supervised SVAE on paired and unpaired datasets and the fingerprint based FC, supervised model on the paired dataset. We tested both models on a test set containing only the molecules from the chosen class. The results are on Figure S1.

For the “Lipids and lipid-like molecules”, semi-supervised SVAE only showed an approximately 15% decrease in the mean fingerprint similarity compared to when the lipids spectra were present (Figure S1A). The more specialised FC, supervised model showed a much larger drop in performance of approximately 55% less correct molecules in the top k candidates (1 <= k <= 20), when the lipids spectra were removed from the training set (Figure S1A). However, for the “Benzenoids” class the models ranked differently: 65% drop in performance for SVAE, and only 35% for FC, supervised. Taken together with the results shown in Figure 3, these results on novel molecule classes demonstrate the bi-model semi-supervised approach shows promise towards generalizing to unseen data better than specialized approaches, but further work is needed to optimize the bi-modal semi-supervised model presented here to the task of compound identification from missing spectra or structures.

Examples of molecules and spectra reconstructions by semi-supervised SVAE from a lipid test set are shown in Figure 5.

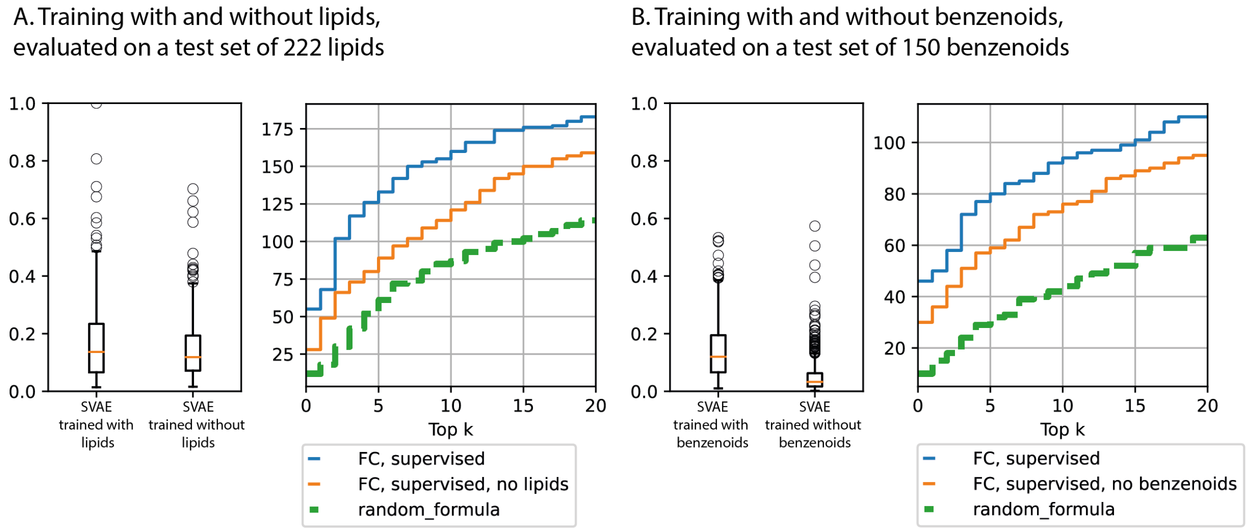


Figure S1: A) Evaluation of spectrum-to-molecule reconstruction quality (fingerprint similarity between the original and the reconstructed molecules) on a test set of 222 “Lipid and lipid-like molecules” for SVAE model and a number of input spectra with the correct molecule in the top k candidates (3-645 candidates for each input spectrum). On a CASMI2017 challenge evaluation this model performed better that SVAE, however after removing the lipid spectra from the dataset the predictive performance for the lipids drops more than two-fold, while the SVAE performance only drops around 15%. B) The same analysis on the “Benzenoids” class shows 65% drop in performance for SVAE, and only 35% for FC, supervised. Different models perform better on different molecule classes.

### C Spectra encoder architecture and dataset

Here we describe how the spectra encoder architecture and a dataset choice were made. The experiments were done using the spectra-to-fingerprints model where a structure fingerprint is predicted from a spectrum. CASMI2017 was used for testing the performance.

Fragmentation tree is an object calculated from a mass spectrum with SIRIUS software (Böcker and Dührkop, 2016). It predicts the most probable chemical formula of a compound and subformulas that correspond to different peaks organised in a tree structure. A child in a fragmentation tree is a subformula that was a part of a parent formula. The fragmentation tree can be seen as an orthogonal source of information for the deep neural network. To process a tree-like structure we used TreeLSTM (Tai et al., 2015). TreeLSTM module output is then concatenated with one of the hidden layers of the multilayer perceptron that processes spectra.

We tested two possible spectra encoder architectures on the two datasets. *Merged* dataset contains the spectra, where all the collision energies for the same InChi Key are merged into one. *Merged+unmerged* dataset has the original MoNA spectra added to the merged dataset. For testing the TreeLSTM architecture, fragmentation trees were calculated for both merged and unmerged spectra.

The results are presented at the Figure S2. Using the unmerged spectra gave the biggest boost in performance, setting the algorithm to the 2nd place among the CASMI2017 contestants. While fragmentation trees improve the performance for the merged dataset, the performance for both architectures is the same if trained on the merged+unmerged dataset. For the purpose of the current study we chose the multilayer perceptron as the spectra encoder architecture and the merged+unmerged dataset. Future work will include further investigation of spectra encoder architectures.

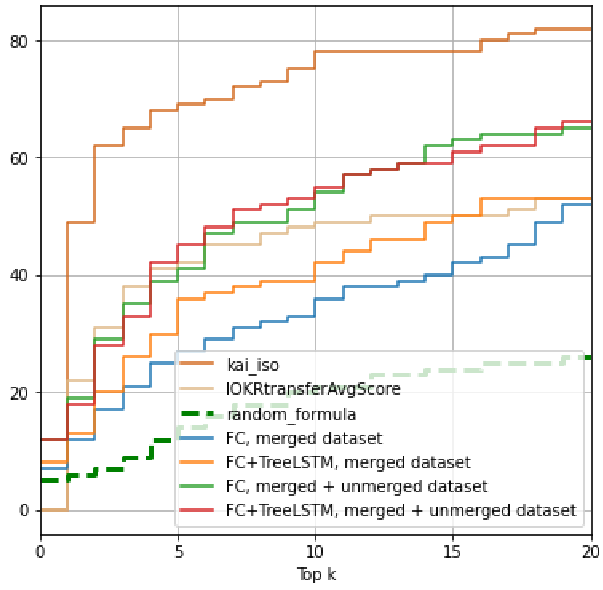


Figure S2: CASMI2017 leaderboard. The x-axis is a number of top candidates, the y-axis shows for how many of 112 positive ion mode spectra the correct molecule showed up in top $k$ candidates. *kai_iso* and *IOKRtransferAvgScore* are the top performant competing algorithms. *FC* is an architecture S1 with only the fully connected layers. *FC+TreeLSTM* is an architecture with fully connected layers and TreeLSTM. *merged* dataset contains only the merged spectra, *merged+unmerged* dataset has merged spectra and unmerged MoNA spectra.

#### D Adding unpaired spectra

Because using the unpaired molecules for training the SVAE model was beneficial, we want to test if the unpaired spectra should be used as well. Additional positive ion mode MoNA spectra were used as the unpaired samples, the list of ids can be found at `https://github.com/sgalkina/svae_spectra`. We ran this experiment for both merged and unmerged spectra.

The Figure S3 shows that using unpaired spectra made the SVAE performance worse. One explanation for this is that the spectra come from a different instrument, thus from a different data distribution, and the resulting latent space becomes harder to model as a Gaussian. However, this hypothesis requires further investigation. We did not use unpaired spectra in this study.

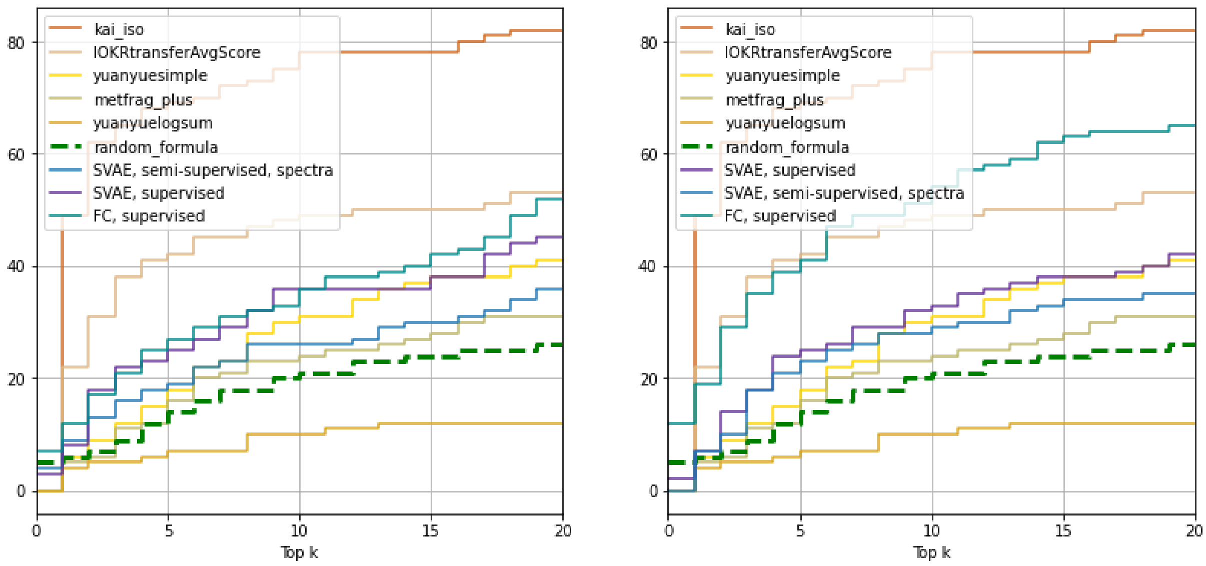


Figure S3: CASMI2017 leaderboard. The x-axis is a number of top candidates, the y-axis shows for how many of 112 positive ion mode spectra the correct molecule showed up in top $k$ candidates. *kai_iso*, *IOKRtransferAvgScore*, *yuanyuesimple*, *metfrag_plus* and *yuanyuelogsum* are the competing algorithms. *FC, supervised* is a deep learning network that predicts a structural fingerprint from a spectrum. *SVAE, semi-supervised, spectra* is trained with the unpaired spectra.

#### E Encoders and decoders network architectures

The encoder and decoder architectures for each experiment and modality are listed below. Adam optimiser is used in all the models. The training algorithm for semi-supervised models iterates over the paired samples around 50 times during one pass through the unpaired molecules set. It means that during $n$ epochs of a semi-supervised model it is exposed to a supervised set for roughly $50n$ epochs. The networks are trained until convergence (no further improvement in aggregated similarity scores for the validation set): 30 epochs for semi-supervised SVAE, 200 epochs for SVAE, 150 epochs for JMVAE, 150 epochs for *FC, supervised*. The latent space dimension is 300.

The fingerprint that was used for spectra-fingerprints predictions is a concatenated vector of Klekota-Roth (Klekota and Roth, 2008), MACCS and PubChem (CACTVS) fingerprints that has 2149 bits in total after removing the duplicated keys.

| ***FC, supervised*** |
| --- |
| Input $\in \mathbb{R}^{11000}$ |
| FC. 10000 & BatchNorm1d & ReLU |
| FC. 5000 & BatchNorm1d & ReLU |
| FC. 3000 & ReLU |
| FC. 2149 & Sigmoid |

Table S1: Network architecture of *FC, supervised*

| **Encoder** |
| --- |
| Input $\in \mathbb{R}^{11000}$ |
| FC. 10000 & BatchNorm1d & ReLU |
| FC. 5000 & BatchNorm1d & ReLU |
| FC. 2048 & BatchNorm1d & ReLU |
| FC. $L$, FC. $L$ |

| **Decoder** |
| --- |
| Input $\in \mathbb{R}^{L}$ |
| FC. 5000 & BatchNorm1d & ReLU |
| FC. 10000 & BatchNorm1d & ReLU |
| FC. 11000 & clamp(0, 100) |

Table S2: Network architectures for spectra encoders and decoders.

| **Encoder** |
| --- |
| Input $\in \mathbb{R}^{250x67}$ |
| 9x9 conv1d 9 & ReLU |
| 9x9 conv1d 9 & ReLU |
| 11x11 conv1d 10 & ReLU |
| FC. 870 & SeLU |
| FC. $L$, FC. $L$ |

| **Decoder** |
| --- |
| Input $\in \mathbb{R}^{L}$ |
| FC. $L$ & SeLU |
| GRU($L$, 488, 3) |
| FC. 67 & Softmax |

Table S3: Network architectures for SMILES strings.

| **Encoder** |
| --- |
| Input $\in \mathbb{R}^{2149}$ |
| FC. 1024 & ReLU |
| FC. 1000 & ReLU |
| FC. $L$, FC. $L$ |

| **Decoder** |
| --- |
| Input $\in \mathbb{R}^{L}$ |
| FC. 500 & ReLU |
| FC. 1024 & ReLU |
| FC. 2149 & sigmoid |

Table S4: Network architectures for fingerprints.

### F Qualitative examples

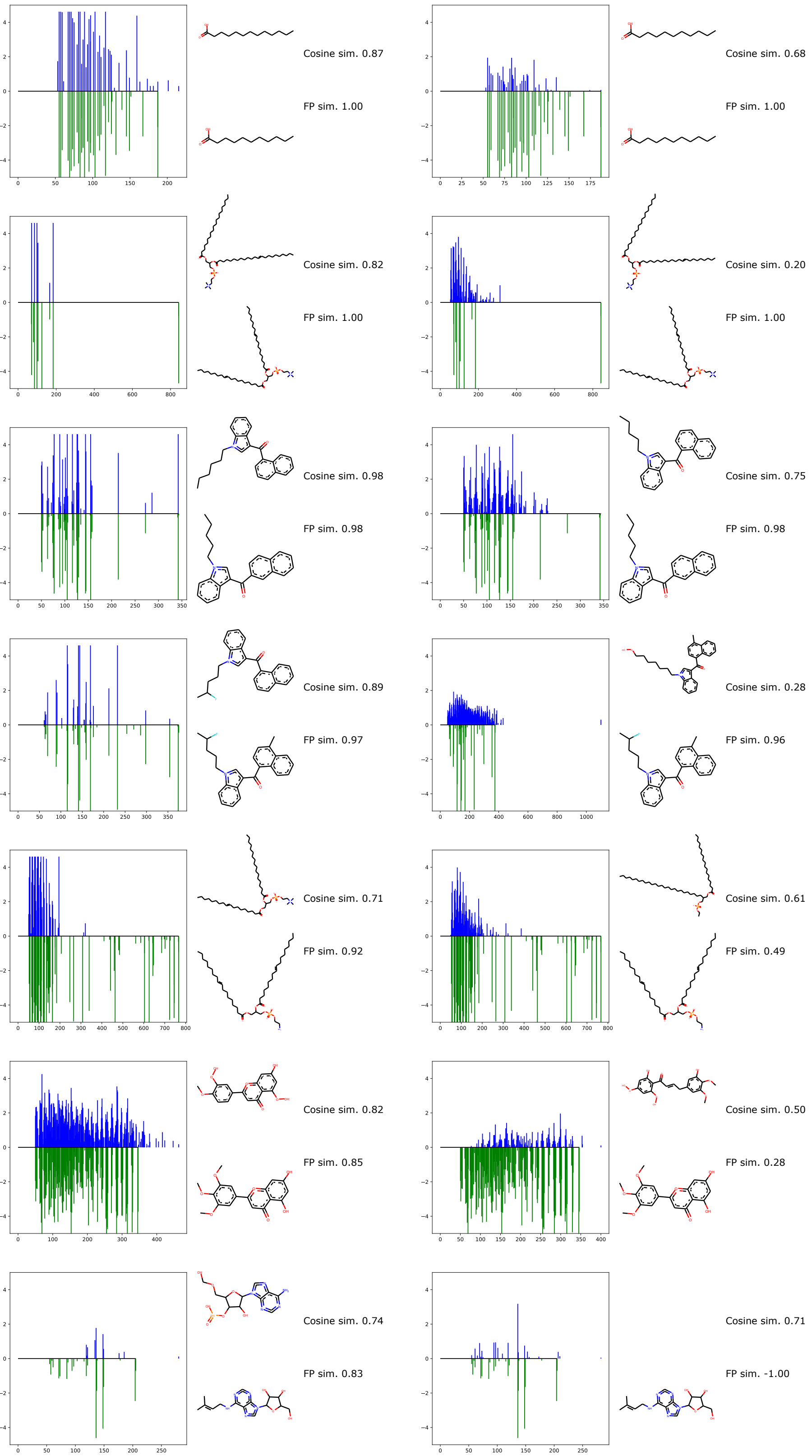


Figure S4: SVAE, semi-supervised, the samples with top 7 fingerprint similarities.

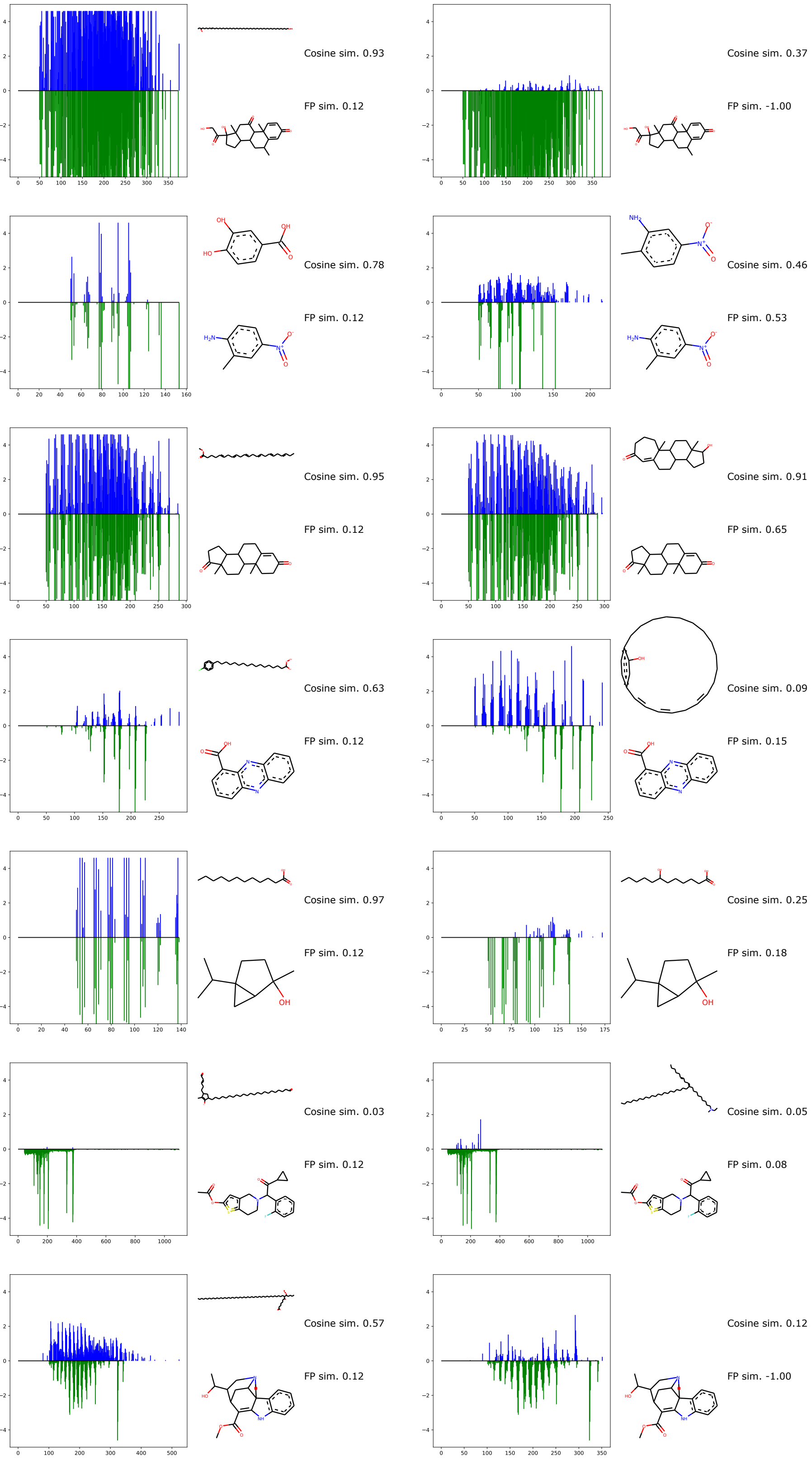


Figure S5: SVAE, semi-supervised, median fingerprint similarity (indices 495-502 from the sorted list).

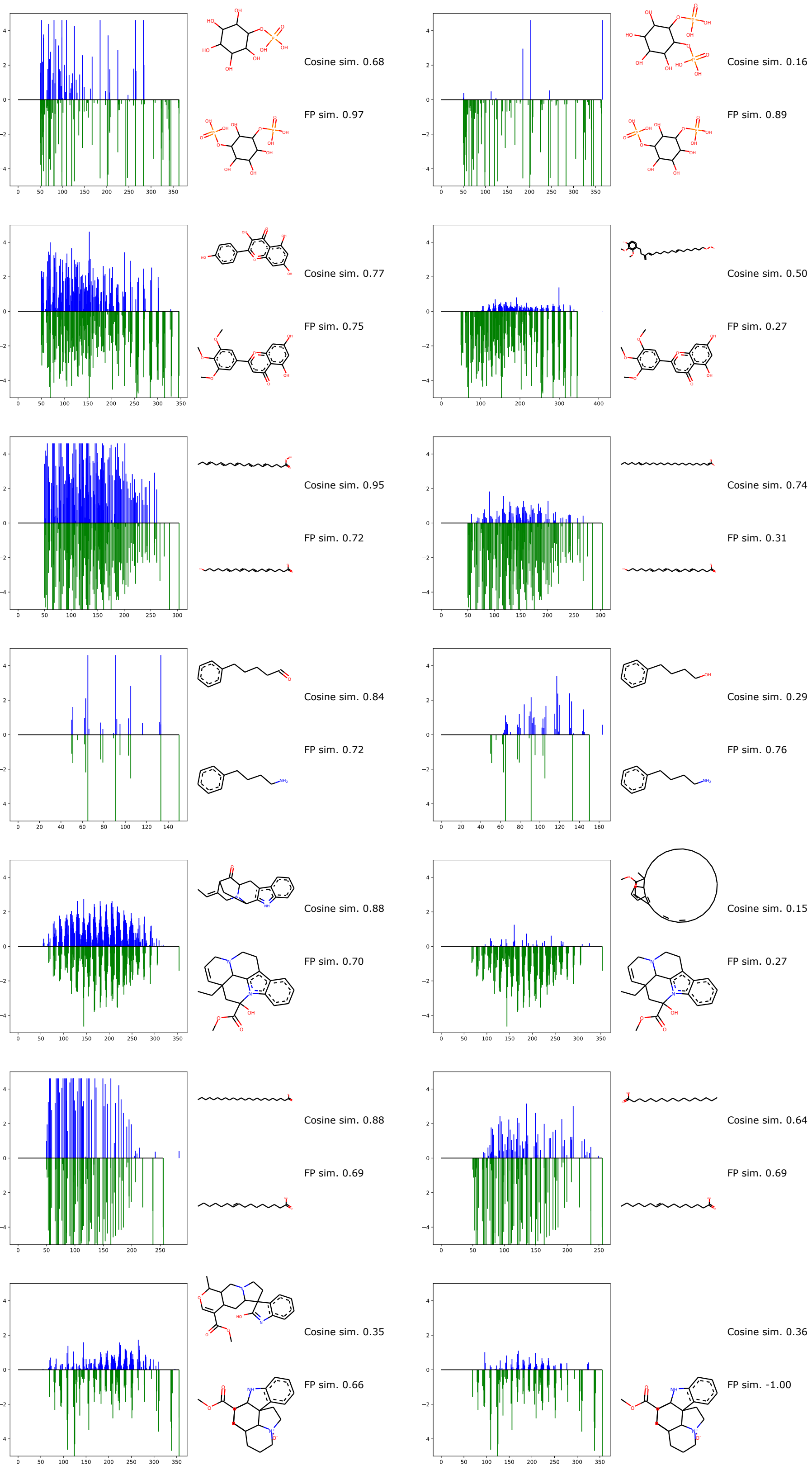


Figure S6: SVAE, supervised, the samples with top 7 fingerprint similarities.

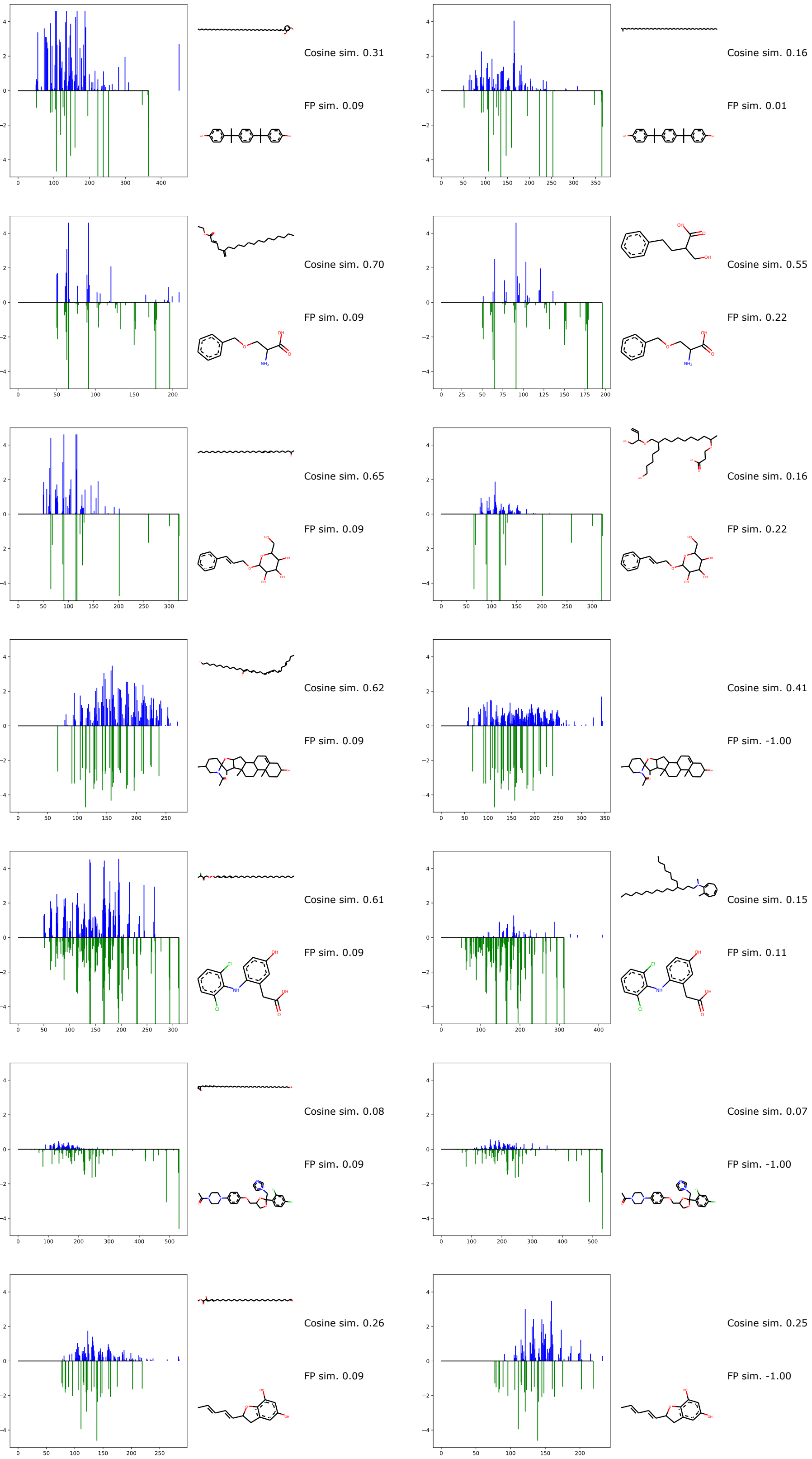


Figure S7: SVAE, supervised, median fingerprint similarity (indices 495-502 from the sorted list).

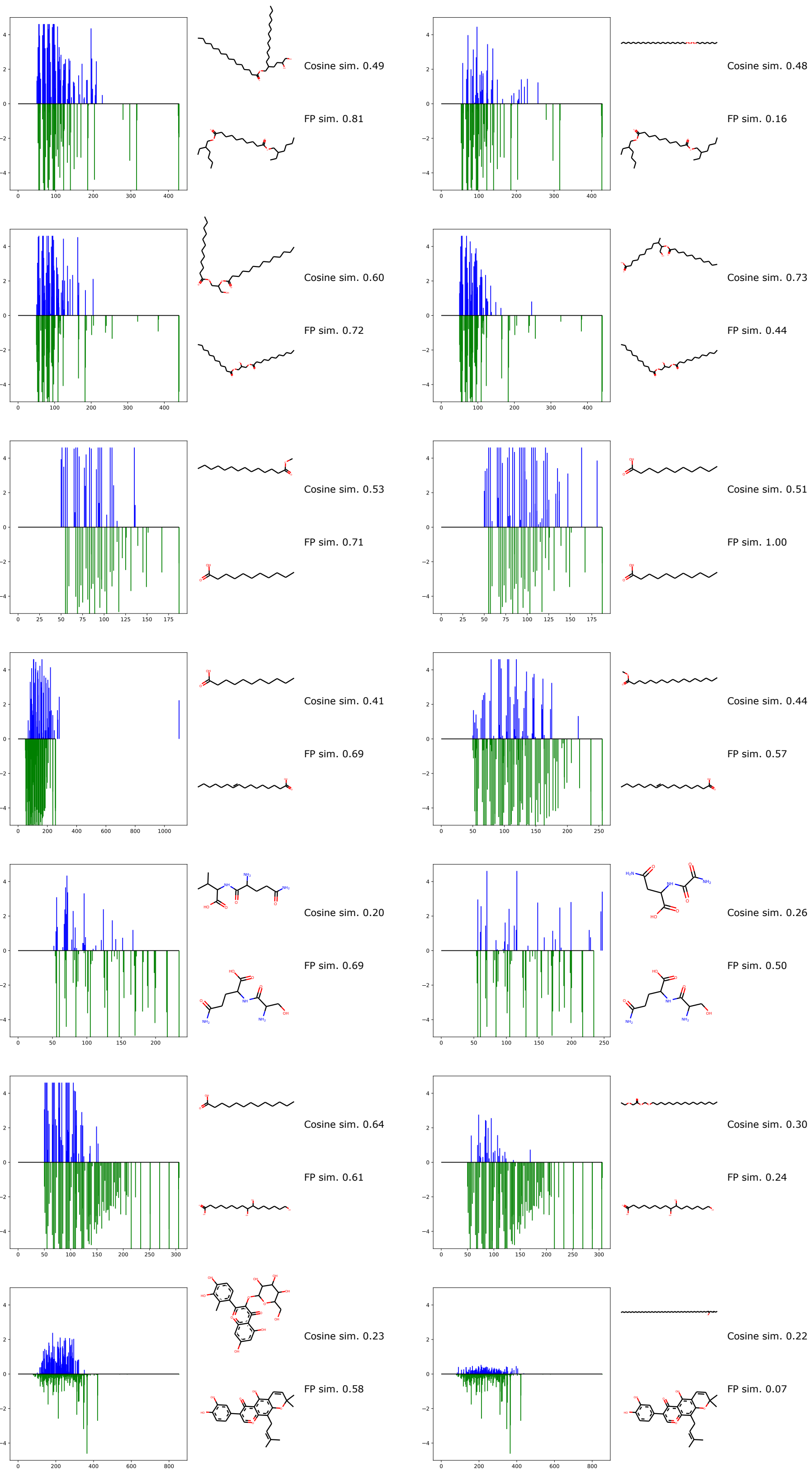


Figure S8: SVAE, supervised, the samples with top 7 fingerprint similarities.

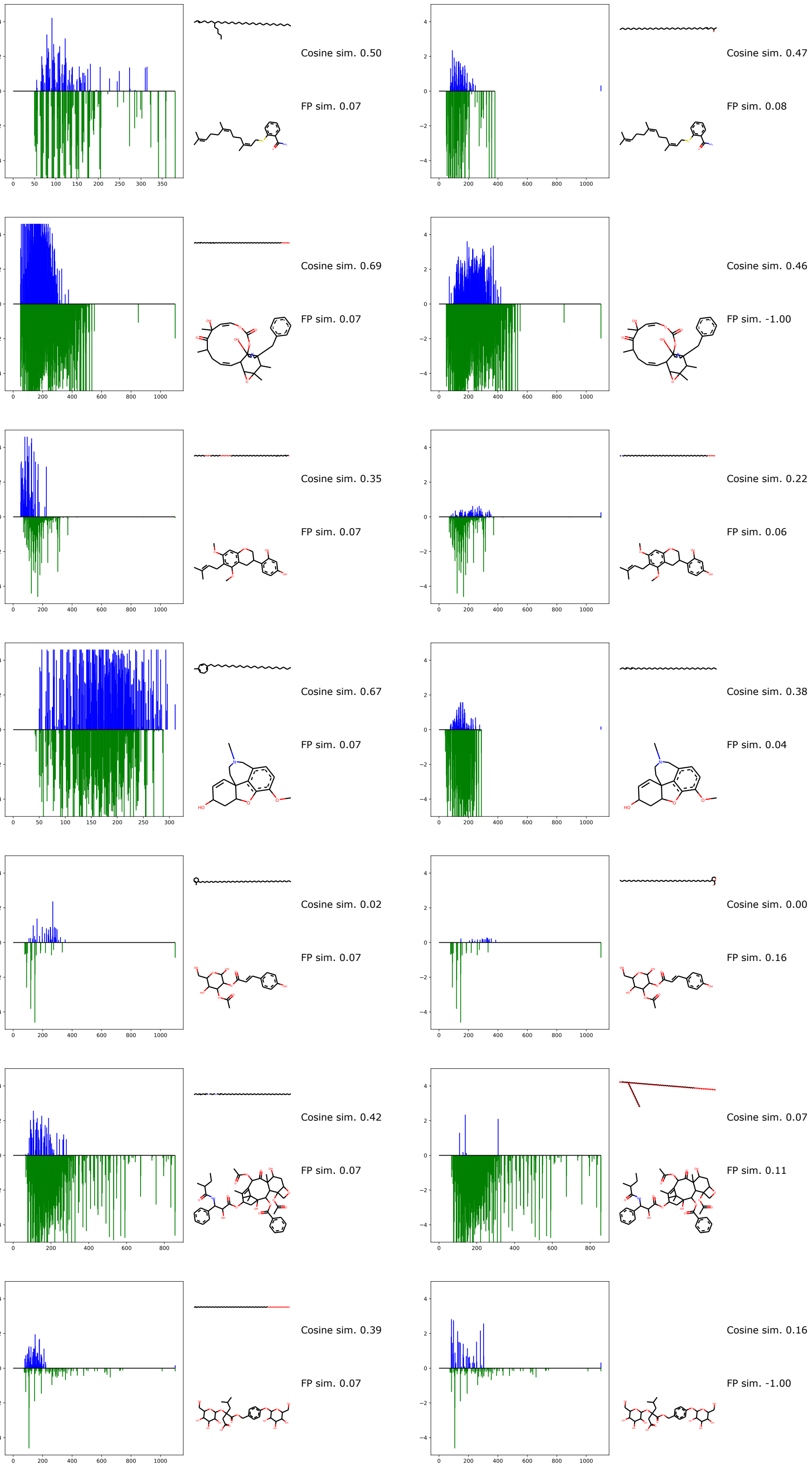


Figure S9: SVAE, supervised, median fingerprint similarity (indices 495-502 from the sorted list).
